# seq-seq-pan: Building a computational pan-genome data structure on whole genome alignment

**DOI:** 10.1101/188904

**Authors:** Christine Jandrasits, Piotr W Dabrowski, Stephan Fuchs, Bernhard Y Renard

## Abstract

**Background:** The increasing application of next generation sequencing technologies has led to the availability of thousands of reference genomes, often providing multiple genomes for the same or closely related species. The current approach to represent a species or a population with a single reference sequence and a set of variations cannot represent their full diversity and introduces bias towards the chosen reference. There is a need for the representation of multiple sequences in a composite way that is compatible with existing data sources for annotation and suitable for established sequence analysis methods. At the same time, this representation needs to be easily accessible and extendable to account for the constant change of available genomes.

**Results:** We introduce seq-seq-pan, a framework that provides methods for adding or removing new genomes from a set of aligned genomes and uses these to construct a whole genome alignment. Throughout the sequential workflow the alignment is optimized for generating a representative linear presentation of the aligned set of genomes, that enables its usage for annotation and in downstream analyses.

**Conclusions:** By providing dynamic updates and optimized processing, our approach enables the usage of whole genome alignment in the field of pan-genomics. In addition, the sequential workflow can be used as a fast alternative to existing whole genome aligners. seq-seq-pan is freely available at https://gitlab.com/groups/rki_bioinformatics

## Background

Thanks to the continuous advances in next generation sequencing (NGS) technologies the number of sequenced whole genomes is also continuously increasing. This has led to a 10,000 fold increase in available bacterial genomes over the past 20 years ([1]). As a result complete sequence information for many species and phylogenetic clades has become available. The current approach to handle the diversity of sequences within a single population is to define a single reference genome with an accompanying comprehensive catalog of variants and other variable genome elements present within that population ([2]). Unfortunately, this representation is limited, as complex genetic differences such as large deletions, insertions or rearrangements cannot easily be expressed in relation to a single reference genome ([3]). This presents a significant drawback, since a combined representation of all genomic content of a species or population that captures the full information on similarity and variation between individual genomes is essential ([4]). Therefore, the more versatile concept of using multiple instead of a single reference genome for common analyses of NGS data is attracting more and more attention.

Initially defined to be the sum of core and dispensable genes of all strains of one bacterial organism ([5]), the term pan-genome is now more commonly used to describe any set of associated sequences aiming for a collective analysis. Gathered under a newly evolving field termed computational pan-genomics, several methods for the generation of data structures that can represent a set of multiple sequences have been developed. These data structures generally aim to fulfill the following requirements: (i) easy construction and maintenance, (ii) adding and retrieving of (biological) information, (iii) comparison to other sets of genomes or short or long sequences from individuals, (iv) easy visualization and (v) advanced data storage ([4]).

We assessed a collection of tools applied for the analysis of multiple sequences (Table 1). Many of these tools use graphs to represent the pan-genome and focus on efficiently building and storing that graph ([2, 6–9]]). Some ([10–12]) focus on subsequent analyses such as mapping reads to the pan-genome, while others ([13, 14]) improve variant detection by using a set of reference sequences instead of a single one. The final category in our collection is made up by tools that introduce a complete data structure and provide methods for the construction, storage, processing and visualization of the pan-genome ([3, 13, 15–17]]). Most of these tools depend on information on the (dis-)similarity of genomes from a multiple genome alignment or a reference sequence with an adjoining corresponding set of variants to create a pan-genome. This prerequisite cannot represent structural variants (e.g. large deletions or insertions or rearrangements of sequences) in most cases and has to be obtained via external tools.

**Table 1.**
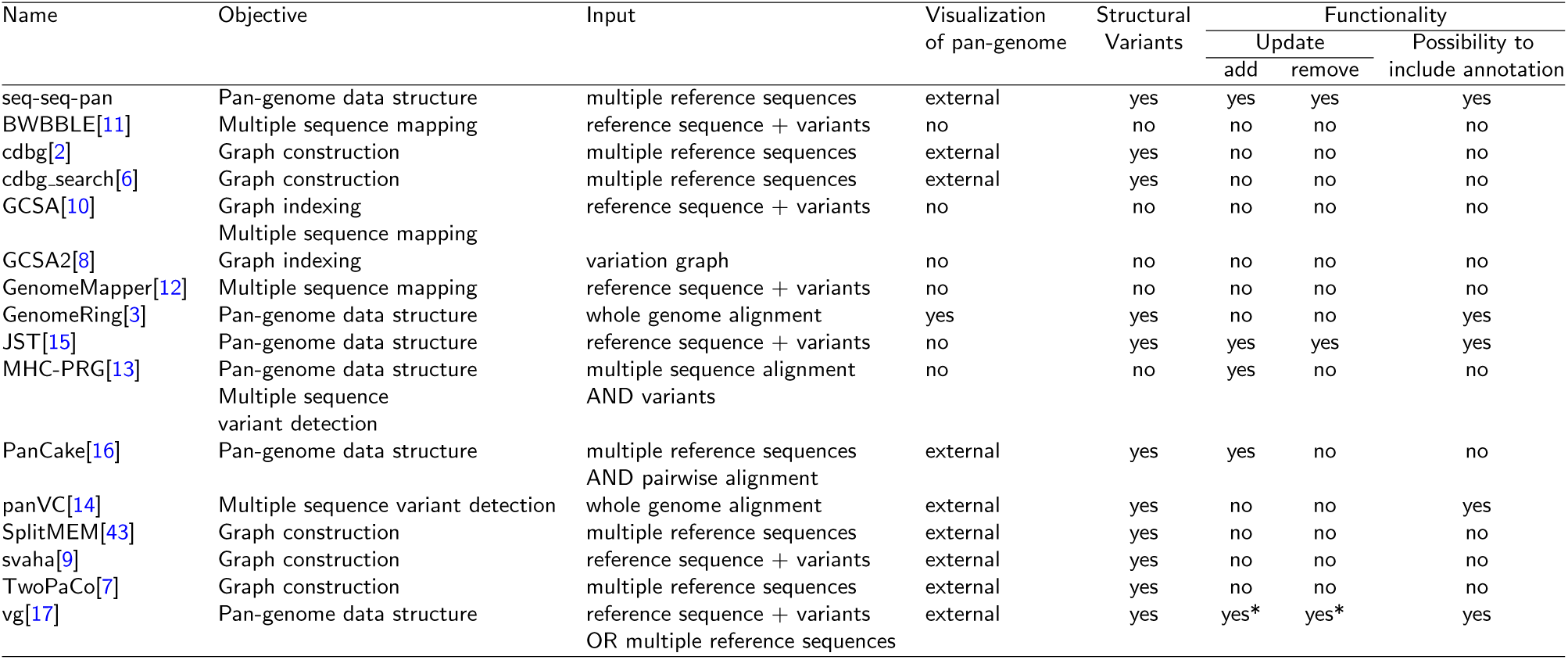
Comparison of pan-genome tools. We analyzed tools for pan-genome analysis that are available or currently under development. This table lists the corresponding publications or websites. We compared the intended use cases of the tools and the prerequisite data required in order to use them. We evaluated the availability of features needed to work with the pan-genome in subsequent analyses, e.g. updating the set of included genomes. Furthermore, we assessed whether the proposed data structures take into account structural variants and whether it is possible to visualize the resulting pan-genome. * Adding and removing of genomes in vg can be achieved using a combination of several steps.

While four of the analyzed tools - JST ([15]), MHC-PRG ([13]), PanCake ([16]), and vg ([17]) - provide methods for adding or removing genomes from the pan-genome data structure, only GenomeRing ([3]), JST ([15]), panVC ([14]) and vg ([17]) offer the ability to annotate biological features. This is often caused by the representation of the pan-genome as graphs, for which there is no standard method providing a coordinate system, which severely complicates the use of existing annotation databases and formats. Proposed strategies for such coordinate systems ([18]) do not meet all preferential criteria (spatiality, readability, and backward compatibility) ([4]). Additionally, new methods for essential analyses such as comparing genetic information of individual samples with a graph of reference sequences have to be developed.

Another (well-established) representation of sets of genomes is their alignment. Whole genome alignments (WGA) implicitly provide a coordinate system that allows the translation between pan-genome and strain genome position, enabling annotation of the alignment with biological features of the individual genomes. Due to the extensive research on whole genome alignment ([19–28]), standard formats (eXtended Multi-FastA ([29])and Multiple Alignment Format ([30])), and methods for processing and visualizing WGA results are available ([3, 31–34]]). In summary, WGA structures presently fulfill most of the desirable properties of a pan-genome, but a severe drawback of existing methods is their final, non-updatable alignment result.

We here present seq-seq-pan, a framework that enables the usage of WGA as a pan-genome data structure. We provide methods for adding additional genomes or removing them from a set of aligned sequences and use them to sequentially align whole genome sequences. Throughout the sequential process we take measures to optimize the resulting whole genome alignment and provide a linear representation that can be used with established methods for subsequent analyses such as read mapping and variant detection.

## 1 Methods

### 1.1 seq-seq-pan Workflow

#### Overview

The key notion of seq-seq-pan is to use and optimize fast, well established whole genome alignment methods to construct a pan-genome from an *a priori* indefinite set of genomes. For this part, we use progressiveMauve ([25]), a fast whole genome aligner that accurately detects large genome rearrangements. The alignment result is comprised of a set of blocks of aligned sequences that are internally free from genome rearrangements (referred to as locally collinear blocks (LCBs)). For each LCB we derive a consensus sequence using the concept of majority vote and combine all sequences with delimiter sequences of long stretches of the character ‘N’. These delimiters are inserted to prevent alignment of sequences over block borders in the following step, because blocks are not consecutive in all genomes. After alignment of the consensus genome with the subsequent genome in the set, all LCBs stretching over block borders are separated. Unaligned sequences of each genome are analyzed again, to align sequences that are considered to be contextually unrelated ([25]). The resulting LCBs with sequences from one or both genomes are joined to the previously aligned blocks. The complete alignment of all genomes is reconstructed from the current and pre-venient alignment (Fig. 1). Optimizing measures are taken throughout the workflow to maintain the synteny of the original genomes and avoid accumulation of short, unrelated sequence blocks. Below we describe all steps of the whole workflow in detail. The consecutive order of execution is depicted in Fig. 2 with a detailed visualization of the alignment of initially unaligned sequences in Fig. 3.

**Figure 1.**
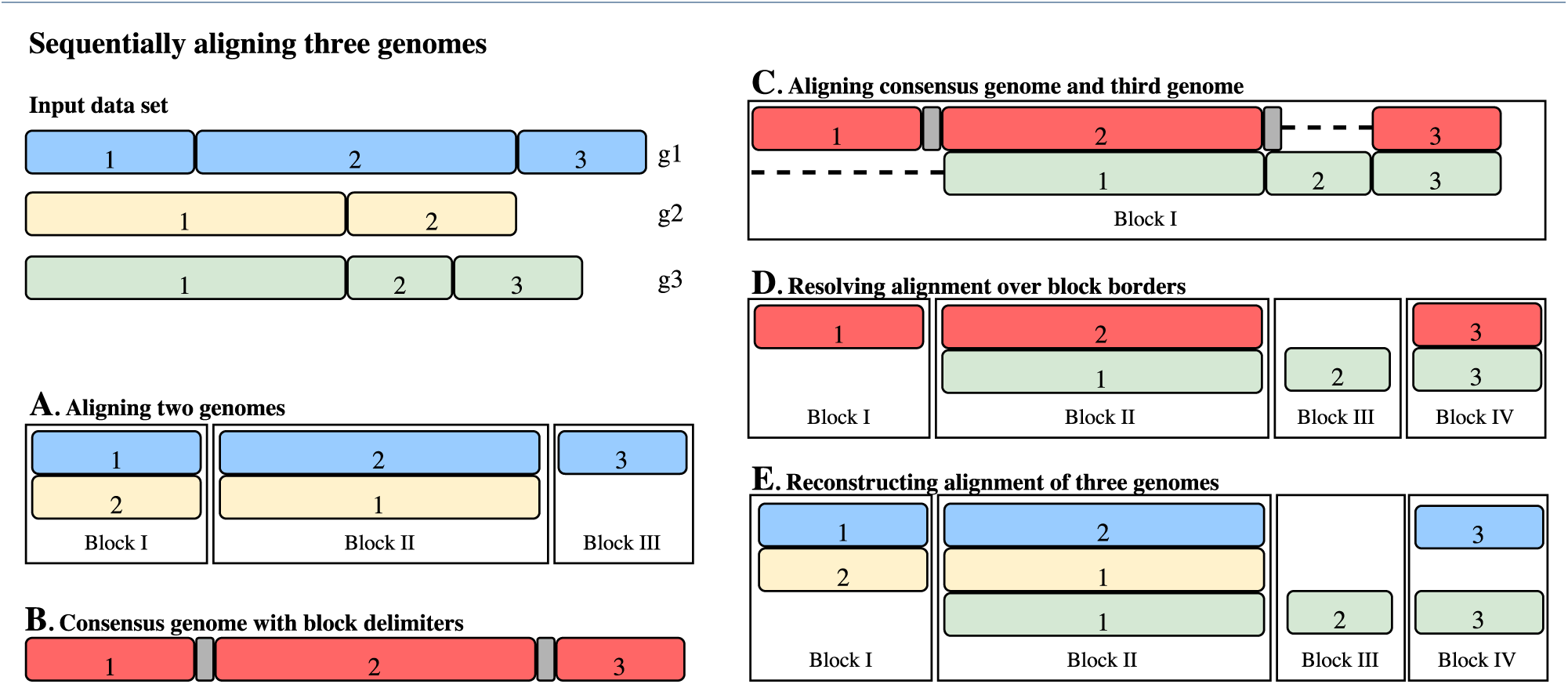
Visualization of the alignment workflow for an example with three genomes. Blocks (blue, yellow, and green) represent whole genomes (g1-3). All sub-sequences are part of locally collinear blocks (LCBs) in the final result. These sub-sequences are marked within the whole genomes and numbered according to their appearance in the respective genome. (A) As a first step, two genomes are selected, aligned and provided as separated blocks of aligned sub-sequences. Block I and II indicate a rearrangement of sub-sequence 1 of g1 when compared to g2. (B) Consensus sequences are built individually for each LCB in the alignment and concatenated with stretches of ‘N’ as delimiters to form a consensus genome (depicted in red with delimiters in gray). (C) The consensus genome is aligned with the third genome (g3), yielding a single LCB. (D) Blocks resulting from alignment with the consensus genome are broken up into smaller blocks at delimiter positions. (E) Alignment of all three genomes is traced back using the newly formed blocks and the alignment of step A. The resulting LCBs can be used as input for step B, and alignment of additional genomes is achieved by consecutive repetition of steps B-E.

**Figure 2.**
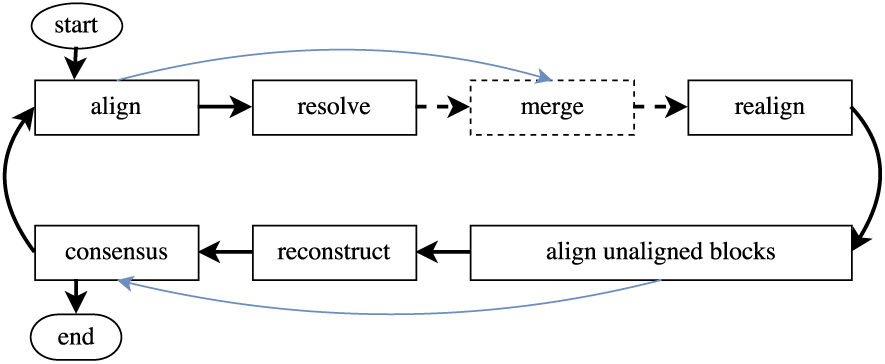
Detailed sequential workflow. Blocks represent steps in the workflow, dashed ones are optional. The first iteration is different and outlined with the blue lines. Subsequent iterations are represented with black arrows. After aligning two genomes - two original ones in the first iteration and a consensus genome with another original one in all following-the result is optimized with an optional merge step and local realignment of sequences around consecutive gap stretches. Initially unaligned blocks are aligned again and resulting blocks are joined with the LCBs containing originally aligned sequences. Then the consensus genome is constructed using the optimized alignment. The consensus genome is aligned with another original genome. Then alignments over block borders are resolved and the alignment is optimized by merging and realignment. Again, initially unaligned blocks are aligned separately. After joining with LCBs of initially unaligned sequences, the full alignment of all genomes is reconstructed.

**Figure 3.**
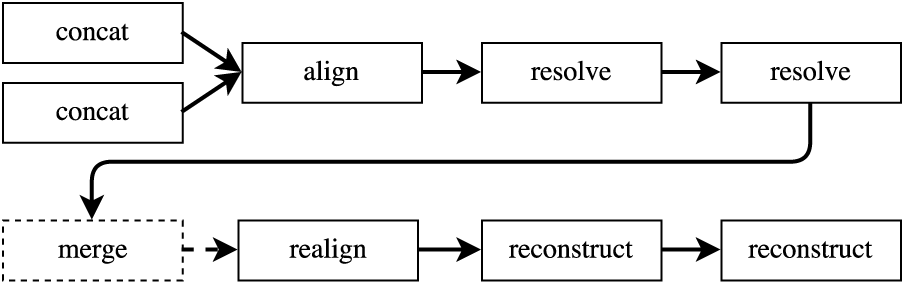
Details of alignment of unaligned blocks. Blocks represent steps in the workflow, dashed ones are optional. Unaligned sequences of each genome are sorted and concatenated with ‘N’ stretches as delimiters. The two constructed sequences are aligned and resulting blocks stretching over borders of concatenated blocks of each genomes are resolved successively. The optional merging step and realignment of sequences around consecutive gaps are followed by the sequential reconstruction of the original sequences of both genomes.

#### Alignment step

When aligning two sequences with progressiveMauve ([25]), the alignment is partitioned in locally collinear blocks to allow for the representation of structural differences such as inversions or translocations. Each resulting block contains parts of either both or one of the genomes locally collinear. They form the basis for the subsequent workflow steps.

#### Merging step

Sequences specific for one of the genomes are also reported as LCBs, but these only contain parts of this one genome. LCBs containing only a single unaligned sequence are typically moved to the end of the alignment file. They are created to ensure collinearity within blocks and can sometimes be of small length. When used in a sequential workflow, it can be advisable to avoid assembling short one-sequence-LCBs and attaching all of them to the end of the consensus genome. We prevent the accumulation of small blocks by merging short one-sequence-LCBs with their neighboring blocks within the genome and realigning consecutive gap stretches (see Realignment step). These short blocks can not only emerge in the alignment step but also result from the resolving step, when a LCB is split at block border positions (see Resolving step).

#### Realignment step

Alignment is improved by realigning genomes at sites where a gap ends in one sequence and starts in the other (referred to as “consecutive gaps”). We scan through the whole alignment and identify all positions with consecutive gaps. Then we extend the interval to the sequence on both sides of the gap sequences by the length of the longer sequence or up to block borders and align these sequences again.

#### Consensus genome construction

LCBs are combined into a consensus genome by concatenating the consensus sequence of each block. At each position within the LCB, all aligned sequences are compared and the most abundant base is chosen for the consensus sequence. In case of ties, the base is drawn randomly from the available choices. To prevent alignments across block borders when the consensus genome is used for alignment, we integrate a sequence of 1000 ‘N’ (undefined nucleic acid) between the consensus sequence blocks into the final sequence. In addition to the consensus genome, two accompanying index files are created. One contains the start positions of all delimiter sequences within the consensus genome and therefore enables the reconstruction of the alignment of all genomes from the alignment of the consensus genome with an additional sequence (referred to as “consensus index file”). The second index file contains the coordinates of all gaps of all sequences per block in the consensus genome, improving the performance of the reconstruction step and the mapping of coordinates between genomes. Furthermore, we make note of the sequence identifier and the description of all genomes and chromosomes, as this information is not contained in the final output of progressiveMauve.

#### Resolving step

Aligning a genome with a consensus genome can result in alignments that span the borders of the LCBs making up the consensus genome. We identify these blocks using the consensus index file. Then, we split them at the start and end of the delimiter sequence. If the alignment spans a complete delimiter sequence the separation results in three new blocks: the first and third one contain the aligned sequences of the two genomes. The second one includes only the sequence of the new genome that was aligned to the delimiter sequence. All gaps contained in this block are removed. In cases where the delimiter sequence is matched with a gap sequence only, we discard the complete block.

#### Alignment of initially unaligned sequences

We take the forward representation of all one-sequence-blocks per genome and sort them. We concatenate the sequences, again integrating stretches of 1000 ‘N’, ending up with one sequence for each of the genomes. These sequences are then aligned using the same process as with the full genomes. Alignment with pro-gressiveMauve, the optional Merging Step and the Realignment Step, are followed by a two-step Resolving and Reconstruction process using each of the initially concatenated sequences as “consensus sequence” (see Fig. 3 and Fig. 1). All blocks with newly aligned and unaligned sequences are joined with the initially aligned blocks for the final Reconstruction step.

#### Reconstruction step

All previous steps result in a set of LCBs that contain parts of the consensus genome, the aligned genome or both. For all LCBs that include consensus genome sequences, we reconstruct the alignment that formed this consensus sequence in the previous workflow iteration. For this we use the coordinate system of the consensus genome and the index file containing delimiter sequence positions. We translate the start and end positions of the consensus sequence in each LCB to their positions within the original genomes. With this information we can extract the bases, gaps and positional information of all sequences and report the complete alignment of all genomes for the current workflow iteration.

#### Removing a genome

After removing a genome from a pan-genome, gaps that were introduced only for the alignment of the removed genome are cut from the remaining genomes. Adjacent LCBs that are now composed of consecutive regions of the same set of genomes are joined to form one LCB.

#### Implementation

For the alignment of pairs of genomes we use progressiveMauve (snapshot 2015-02-13). All other parts of the sequential workflow are implemented in Python3.4. For performance reasons, the consensus construction step was additionally implemented in Java8. We use the following Biopython (version 1.68) modules ([35]) in our workflow: SeqIO, Seq and SeqRecord, for reading, writing and manipulation of single sequences and pairwise2 for aligning two sequences in the Realignment step. For performance reasons we use blat ([36]) for the alignment of large sequences in the Realignment step.

For a straightforward construction of a pan-genome from a set of genomes, we combined all steps with the workflow management software Snakemake ([37]). The pipeline can be parametrized to include the optional merging steps and all necessary steps are determined automatically in each iteration. The output is composed of the final alignment of all input genomes and the corresponding consensus genome including index files necessary for following analyses and further updates of the pan-genome.

### Setup for comparison experiments

#### Data

We use the set of reference genomes of *Mycobacterium tuberculosis* available in the NCBI RefSeq database as of November 30^*th*^, 2016 throughout all experiments. This set contains 43 complete genomes. Accuracy of alignment was tested on a simulated dataset of twelve genomes with the genome of *Escherichia coli* K12 as basis for the simulation of evolution. For evaluating the runtime when adding an additional genome to a pan-genome, we used another *M. tuberculosis* genome that became available on December 26^*th*^, 2016 (for details on genomes see Additional file 1).

#### Simulated Data

Accuracy of alignment was tested on a simulated dataset of 13 genomes. For the simulation of genomes with a known true alignment we used the EVOLVER software ([38]) and the evolverSimControl suite ([39]) as described in the Alignathon project ([40]). The tool evolverSimControl enables the user to simulate several genomes along a phylogeny with EVOLVER. We used an *E. coli* K12 genome (NC 000913.3) as the origin of the evolution simulation. For the evolution parameters we adapted the example provided by the EVOLVER team. We fit the parameters to the smaller size of the *E. coli* genome by changing the probabilities of large insertion and deletion events and setting the maximum size of these events to 7000 – roughly the size of the longest *E. coli* gene. We simulated twelve genomes without using mutation acceptance constraints with the phylogeny depicted in Fig. 4.

**Figure 4.**
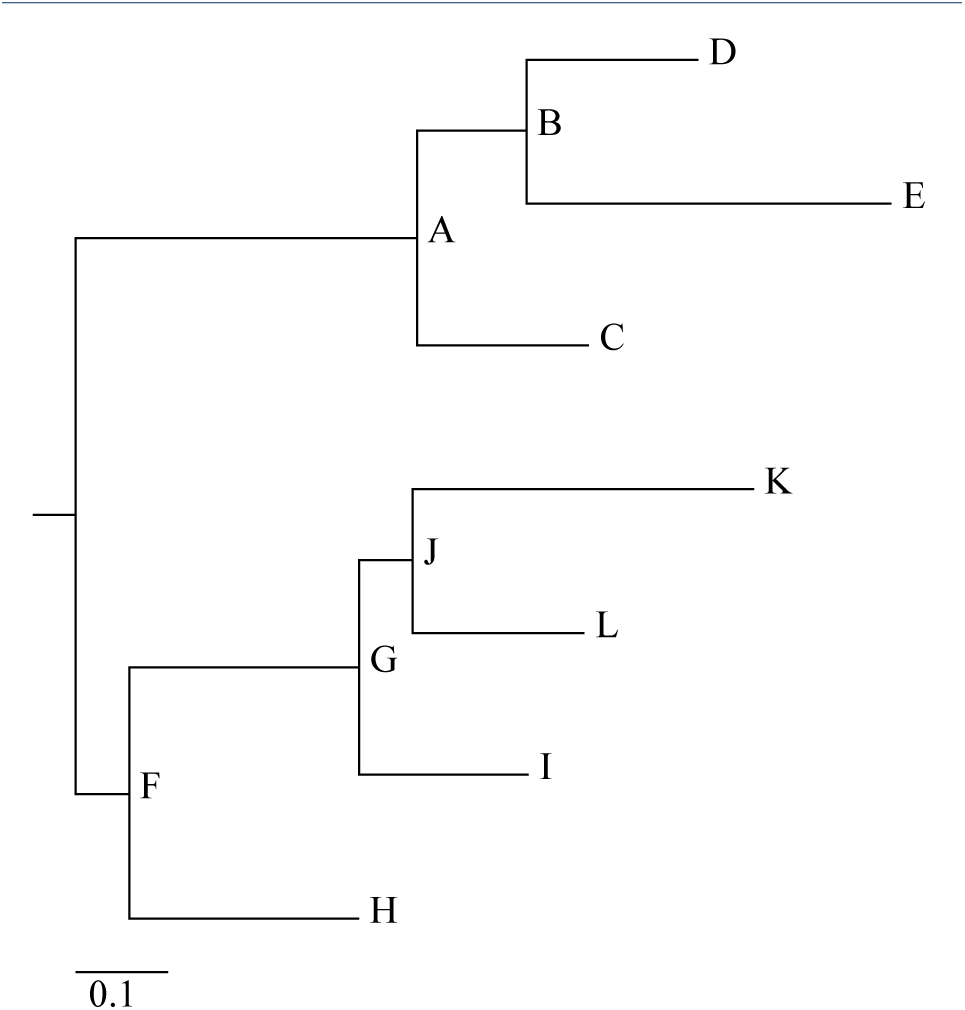
Visualization of the phylogenetic tree used to simulate genomes with EVOLVER. The corresponding NEWICK tree is (((D:0.015625,E:0.0333)B:0.01,C:0.015625)A:0.03125, (((K:0.03125,L:0.015625)J:0.005,I:0.015625)G:0.02083, H:0.02083)F:0.005);. (drawn with online version of Phylodendron ([42]))

#### Comparison of alignments

We use an alignment comparison method to compare our results with the results of other whole-genome aligners: the tool mafComparator from the mafTools collection ([40]). To compare the alignments of the simulated dataset with the true alignment, we calculate recall, precision and F-score as described in the Alignathon project([40]). To use the same method for comparing alignments of the *M. tuberculosis* dataset we choose the alignment of the other aligners to act as the true alignment and our results as the prediction in each comparison. Here, we use the F-score to assess the similarity of the alignments. Depending on the choice of the first genome in the alignment, all sequences in all LCBs are potentially reported on opposite strands when comparing two alignments. In these cases, we replaced all sequences of one alignment with their reverse complement because otherwise mafComparator would report these alignments to be highly dissimilar. Additionally, we compare the length of the whole alignment, and the number of genomes per LCB. With this, we can measure the extent of fragmentation of the alignment and how many bases were aligned.

#### Merging and Sorting

Due to the sequential nature of our workflow, the order in which genomes are added to the pan-genome might influence the resulting alignment. To investigate this effect we arrange the genomes by similarity and by consecutive dissimilarity and compare the results. To sort the input genome sequences by similarity we apply the D2z score ([41]) on all pairs of sequences. The score reflects the sequence similarity, i.e. higher scores stand for more similar sequences. We calculate the upper quartile of all similarity scores and select the genome with the smallest distance from all others. The remaining sequences are ordered by their similarity to this genome.

As an alternative, to obtain a series of strongly differing genomes, we sort them as follows: we again start with the genome with the smallest distance from all others. Then we choose the one with the lowest similarity score as second and the genome most similar to the first for the third position and continue to alternate genomes in this manner throughout the complete set (so the sequence 1, 2, 3, 4, 5, 6 becomes 1, 6, 2, 5, 3, 4). We also compare alignments that were created with and without using the merging steps (See order of genome sets in Additional file 1).

#### Whole genome alignment tools

We compared our sequential genome alignment approach with whole genome alignment tools to review the accuracy of the final alignment. For this, we chose progressiveMauve ([25]), Mugsy ([23]), and progressiveCactus ([19]) as these are commonly used methods that allow aligning non-collinear genomes. Each of these tools separates the final alignment into LCBs. We parametrized all tools to not report duplications and disabled filters on LCB sizes to fit the results to the methology of seq-seq-pan. In addition to comparing the final alignments, we analyze the time and memory needed to create these results. In cases where no ground truth for the alignment is available, we regard the concordance of the results of all tools.

#### Pan-genome tools

For comparison, we choose PanCake, which also accepts whole genomes as input and bases the construction of the pan-genome data structure on sequence alignment methods. PanCake represents all genomic sequences in the form of feature instances. Each feature contains part of a genome sequence and start and stop coordinates within the genome. By using the information of pairwise genome alignments, shared features can be extracted. These features contain a single version of the sequence and a list of edit operations and positional information describing all aligned sequences. Following the recommendations by the authors, nucmer ([24]) was used for pairwise alignments. We measure the time it takes to construct a pan-genome. Tasks that are part of many analyses of pan-genomes include adding an additional genome or removing a genome and extracting a genomic sequence from the pan-genome. Thus, we examine whether these steps are possible and which runtime they require. To account for differences in time needed to extract genomes based on their position within the pan-genome, we extracted each genome once and calculated the average time.

## Results

As we sequentially construct a whole genome alignment, we compare our results with the alignments of progressiveMauve ([25]), Mugsy ([23]) and progressive-Cactus ([19]) for the simulated dataset and the set of 43 *M. tuberculosis* genomes. We compare the runtime and memory requirements of pan-genome construction and the provided functional features between seq-seq-pan and PanCake ([16]) using the *M. tuberculosis* dataset. We show that the order of genomes has minimal effects on the final alignment and that the merging step produces a less fragmented alignment. For all comparison analyses, we show the results for the whole genome alignment constructed with seq-seq-pan from genomes sorted by similarity using the merging step.

### Sorting and Merging

We use two different orders for the simulated and the *M. tuberculosis* dataset and compare the results. The order of sequentially aligned genomes has a minimal effect on the length of the full alignment in all pairs of alignments with and without the merging step (Table 2). We compare the alignments of the simulated dataset with both sort orders to the true alignment with the mafComparator tool from the mafTools suite ([40]). The sort order has minimal effects on the precision and recall (Table 3). Using the sequential workflow without the merging step, results in the alignment of fewer genomes within each LCB in both datasets (Table 2). This indicates that the alignment is more fragmented when small LCBs are not merged to their neighboring blocks. Nevertheless, the F-score comparing the results with and without merging indicates only small differences in the overall alignment (Table 3).

**Table 2.** Effect of sort order. We compare the results from sequentially aligning two genome datasets in two different orders including and excluding the merging step in the workflow. We estimate the fragmentation of the alignment via the total alignment length and the number of sequences per block.

|  |  | Total Alignment Length | Mean number of Sequences in LCB |
| --- | --- | --- | --- |
| Simulated dataset (13 genomes) |  |  |  |
| Similar | with Merging Step | 4809015 | 9.2 |
| Dissimilar | with Merging Step | 4777972 | 7.4 |
| Similar | NO Merging Step | 4789770 | 5.5 |
| Dissimilar | NO Merging Step | 4757489 | 5.0 |
| *M. tuberculosis* dataset (43 genomes) |  |  |  |
| Similar | with Merging Step | 4826979 | 16.1 |
| Dissimilar | with Merging Step | 4842752 | 14.9 |
| Similar | NO Merging Step | 4859842 | 7.5 |
| Dissimilar | NO Merging Step | 4776700 | 7.2 |

**Table 3.**
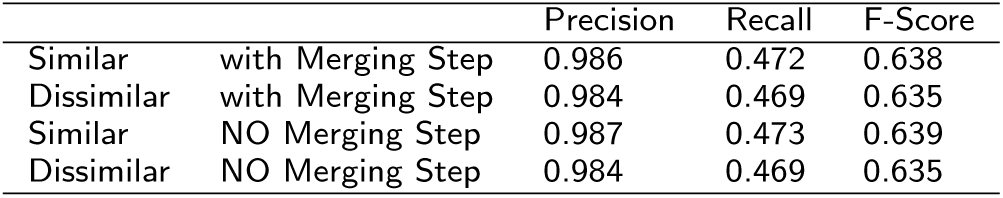
Accuracy of alignment. We compare results of aligning the simulated genomes with seq-seq-pan including and excluding the merging step in the workflow in two different sort orders with the true alignment.

|  |  | Precision | Recall | F-Score |
| --- | --- | --- | --- | --- |
| Similar | with Merging Step | 0.986 | 0.472 | 0.638 |
| Dissimilar | with Merging Step | 0.984 | 0.469 | 0.635 |
| Similar | NO Merging Step | 0.987 | 0.473 | 0.639 |
| Dissimilar | NO Merging Step | 0.984 | 0.469 | 0.635 |

### Comparison with whole genome alignment tools

The results of all whole genome aligners and our approach are competitive. In the simulated setting seq-seq-pan achieves similar precision and recall as pro-gressiveMauve and Mugsy and better results than progressiveCactus (Table 4). We assessed whether the results of progressiveCactus and Mugsy improved when parametrized to detect duplications. This had almost no effect for Mugsy and improved the comparatively low precision for progressiveCactus, but reduced the recall. All aligners achieve low recall, but comparably low values were also observed for simulated datasets used in the publication introducing the comparison method applied here ([40]). For the *M. tuberculosis* dataset, our results are closer to the alignment by progressiveMauve than the ones by Mugsy and progressiveCactus. ProgressiveCactus and Mugsy yield a very similar alignment (Table 5). Table 6 shows the high speed up seq-seq-pan achieves compared to the whole genome alignment tools. Seq-seq-pan aligns 13 simulated genomes within 30 minutes and 43 *M. tuberculosis* genomes within two hours - being at least five times faster than all other tools with the real data set. ProgressiveCactus required almost two days for the alignment of 43 genomes and we were unable to align the whole set with Mugsy. It took Mugsy almost 15 hours to align 39 (randomly chosen) genomes. The memory requirements during the alignment construction are correlated with the elapsed time in most cases and are therefore lowest for seq-seq-pan.

**Table 4.** Precision, Recall and F-Score for alignments of the simulated dataset. We compare the results of all alignment tools with the true alignment of the simulated genomes.

|  | Precision | Recall | F-score |
| --- | --- | --- | --- |
| seq-seq-pan | 0.986 | 0.472 | 0.638 |
| progressiveMauve | 0.984 | 0.473 | 0.639 |
| Mugsy | 0.999 | 0.474 | 0.643 |
| progressiveCactus | 0.892 | 0.473 | 0.618 |
| Mugsy with duplications | 0.999 | 0.474 | 0.643 |
| progressiveCactus with duplications | 0.999 | 0.339 | 0.506 |

**Table 5.** F-score for pairwise comparison of alignment results for the *M. tuberculosis* dataset. We estimate the similarity of alignments of progressiveMauve, Mugsy, progressiveCactus, and seq-seq-pan by calculating the pairwise F-score. * Aligning 43 *M. tuberculosis* genomes caused a segmentation fault in Mugsy. We were able to align 39 genomes and therefore compare the results only for this set of sequences.

|  | seq-seq-pan | progressiveMauve | Mugsy* | progressiveCactus |
| --- | --- | --- | --- | --- |
| seq-seq-pan | - | 0.816 | 0.804 | 0.775 |
| progressiveMauve | 0.816 | - | 0.798 | 0.770 |
| Mugsy* | 0.804 | 0.798 | - | 0.934 |
| progressiveCactus | 0.775 | 0.770 | 0.934 | - |

**Table 6.**
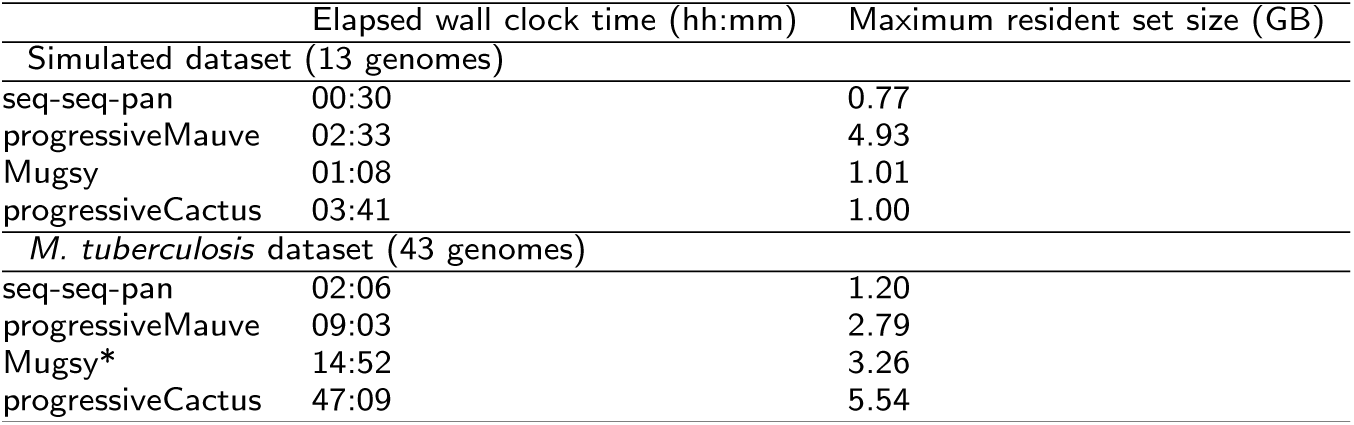
Runtime and memory usage. We compare seq-seq-pan to other whole genome aligners in terms of runtime and memory usage. * Aligning 43 *M. tuberculosis* genomes caused a segmentation fault in Mugsy. This table lists data for aligning 39 genomes with Mugsy, but the whole set of 43 genomes for all other tools.

### Comparison with pan-genome tools

In addition to the set of reference genomes, PanCake requires pairwise alignments of all genomes to construct a pan-genome. In the case of our experiments with 43 *M. tuberculosis* genomes, the construction of 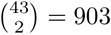 pairwise alignments is required. For our comparison, we calculated these sequentially, but depending on the available hardware, this task can easily be parallelized. For this reason, we list the runtime and memory requirements of pairwise alignments with nucmer ([24]) separately (Table 7). Constructing the pan-genome with PanCake takes considerably longer than with seq-seq-pan. Also, the extraction of genomes or intervals of genomic sequences takes more time. The resulting pan-genome file from PanCake is smaller in size than the one created with seq-seq-pan. The reason for this difference in size and sequence extraction times is the strategy of PanCake of storing only the differences to a reference genome instead of the whole sequence for all genomes within a shared feature. Removing genomes and the generation of a consensus genome are features that are only provided by seq-seq-pan as listed in Table 1.

**Table 7.** Comparison of seq-seq-pan and PanCake. First we compare the runtime and memory usage of pan-genome creation for the set of 43 genomes. We also evaluate the file size of the resulting pan-genome. We clock all available features (adding a genome, extracting part of a genome or the whole genome, remove a genome and constructing a consensus genome). * Extraction times for whole genomes and parts of sequences are equal. We extracted the interval 500-1000 for all genomes. ** Each of the 43 genomes was removed from the whole set.

|  | seq-seq-pan | PanCake | nucmer |
| --- | --- | --- | --- |
| Time for construction (hh:mm:ss) | 02:06:00 | 88:10:00 | 03:04:00 |
| Maximum memory usage | 1.20 GB | 2.34 GB | 0.10 GB |
| Pan-genome file size | 198 MB | 36 MB | - |
| Time to add genome | 00:04:01 | 05:33:52 | 00:08:48 |
| Mean time for extraction of sequence* | 00:00:09 | 00:01:08 | - |
| Mean time for removing genome** | 00:00:19 | not available | - |
| Time for consensus genome creation | 00:00:47 | not available | - |

## Conclusions

In this contribution, we introduced seq-seq-pan which enhances whole genome alignments by adding critical features for pan-genome data structures e.g. updating the set of genomes within the pan-genome. It provides a fast and simple construction process for whole genome alignments while optimizing the results for usage in subsequent analyses. The continuous merging of small unaligned blocks prevents the accumulation of sequences without context or position within the alignment and preserves the synteny of the original genomes, while the realignment of pairwise alignments avoids the introduction of additional repeats into the linear pan-genome representation. Both steps support the application of mapping based methods such as read alignment.

The whole genome alignment format that we use as representation of a pan-genome in seq-seq-pan retains the full sequences and gaps for all aligned genomes in addition to meta-information about block borders. Therefore, it is not suitable to store the pan-genome efficiently. However, this format ensures loss-less and faster handling of the data. Further, it is thereby accessible by currently available downstream analysis tools without requiring subsequent novel tool implementations.

We demonstrate that the sort order of genomes does not substantially influence the result despite the sequential nature of our approach.

We compared seq-seq-pan with three whole genome aligners that offer alignment of non-collinear sequences. These tools use sophisticated methods for the identification of ortho-and even paralogs and conserved sequences. With these features, they identify similar but unrelated sequences within genomes, an aspect that is not considered in the field of pan-genomics. As we do not take such measures, we did not expect very high concordance between our results and the whole genome alignments. However, our comparison shows that our alignment differs as much from the results of progressiveMauve, progressiveCactus and Mugsy as their results differ among each other. Our approach is able to align a set of genomes much faster and with less memory usage than these whole genome alignment tools. Due to the focus on highly conserved sequences, some of these tools also provide a very fragmented alignment with many small blocks, which is prevented by the merge step in seq-seq-pan.

We compare our approach with currently available methods in terms of applicability and needed prerequisites (input data). For a detailed comparison, we chose PanCake as an approach by which a pan-genome can be constructed from a large set of genomes. We show that the construction of the pan-genome and using the structure for basic tasks requires substantially less time with seq-seq-pan than with PanCake. Some features, such as removing a genome from the pan-genome and the construction of a linear presentation of the pan-genome in the form of a consensus sequence, are not directly available in any other pan-genomics tool. For instance, the authors of PanCake focused on the analysis of core and accessory gene sets and therefore provide different functionalities.

In the time between November 30th, 2016 and January 20th, 2017 eight new *M. tuberculosis* genomes became available in the NCBI Ref-Seq database. This already highlights the importance of having the ability to extend a pan-genome structure. Methods such as the investigated whole genome alignment tools that constrain the user to start the alignment afresh with the increased number of genomes are at risk of reaching computational limits (some indications could be observed for Mugsy in the experiments already) which is mitigated by our iterative approach which quickly adds new sequences without having to rebuild previously calculated results. Furthermore, publicly available sets of genomes, such as the collection of “Complete Genomes” in the NCBI RefSeq database, are subject to change due to altered quality standards or the redefinition of reference genomes, such as the commonly used *M. tuberculosis* H37Rv strain. Therefore, it is essential that pan-genome representations also provide the feature to easily remove genomes from the initial set without impacting the remaining genomes. Most of the evaluated tools do not provide methods for updating a constructed pan-genome. Particularly research like molecular surveillance, where new data is continuously analyzed and incorporated, depends on data structures that allow the integration of an up-to-date set of genomes. In summary, we present a data structure for the representation of pan-genomes that provides a unique set of features needed for efficiently working with collections of similar sequences and that can be integrated with existing methods for visualization and subsequent analyses.

## Abbreviations

NGS: next generation sequencing
WGA: whole genome alignment
LCB: locally collinear block

## Declarations

Availability of data and materials

Complete genomes are available from the NCBI RefSeq database. Detailed information on accession numbers are available in Additional file 1. Simulated data is available from the project repository. seq-seq-pan is written in Python and Java and runs on Linux with the following requirements: Java 8, Python3.4 including Biopython 1.68, blat, progressiveMauve, snakemake. It is distributed under the FreeBSD License and available at https://gitlab.com/rki_bioinformatics.

## Author’s contributions

CJ, PWD, FS and BYR designed the workflow and experiments. CJ and PWD wrote the software. CJ executed the experiments. All authors contributed to preparation of the manuscript. All authors reviewed and commented on the manuscript. All authors read and approved the final manuscript.

## Acknowledgements

We would like to thank Lena Fiebig and Walter Haas (Robert Koch Institute) for fruitful discussions and André Hennig and Kay Nieselt (University of Tübingen) for valuable feedback and insight.

## Ethics approval and consent to participate

Not applicable.

## Consent to publish

Not applicable.

## Additional Files

### Additional file 1

(XLSX) – Supplementary Tables for “seq-seq-pan: Building a computational pan-genome data structure on whole genome alignment”

Additional File 1 provides detailed description of the used datasets, including accession numbers and sort order.

